# Artemisinin-Conjugated Phototheranostic Probe for Golgi-Targeted H_2_O_2_ Detection and Side Effect Attenuation

**DOI:** 10.64898/2025.12.02.691769

**Authors:** Jun Liu, Yifen Yang, Zhengqin Liu, Yanli Long

**Affiliations:** School of Chinese Ethnic Medicine, Guizhou Minzu University, Guiyang, 550025, China

**Keywords:** Artemisinin, theranostic probes, side effects, MIDA boronates, H_2_O_2_

## Abstract

Understanding the mechanism by which Artemisinin conjugated with phototheranostic probes for reducing side effects (AC-Pr-RSE) will guide endeavor to improve the therapeutic outcomes. Herein, the first report describes the application of a cytotoxic artemisinin derivative and N-methyliminodiacetic acid (MIDA) boronates conjugated with a naphthalimide fluorophore for the detection of H_2_O_2_ in the Golgi apparatus. A comparative analysis of the localization signals from the fluorescent artemisinin derivative and organelle-specific dyes uncovered that the Golgi apparatus functions as the primary site of its accumulation, which could efficiently enhance the intracellular reactive oxygen species (ROS) level and induce cell apoptosis. This work highlights the potential of AC-Pr-RSE study, which enables us to optimize the design of theranostic probes to improve their biological activities during oxidative stress.

## 1. Introduction

In tropical and subtropical regions, malaria persists as a crucial public health concern.^[1,2]^ While the concerning surge in resistance to antimalarial drugs is progressing at an astonishing pace, diverse antimalarial drugs remain capable of curing malaria.^[3,4]^ Artemisinin (Qinghaosu), a sesquiterpene lactone detected by Nobel Prize winner You-You Tu, has been commonly applied in traditional Chinese medicine against the malarial parasite.^[5]^ In addition to its potent antimalarial activity, emerging studies indicate that artemisinin and its derivatives may possess antifungal, antiviral, anthelmintic and anticancer properties.^[6-8]^ At present, the most reported artemisinin dependent endoperoxide bioactivation take place in the mitochondria, which concurrently triggers the synthesis of ROS (especially H_2_O_2_) and oxidative stress.^[9-11]^ Moreover, a significant similarity was found for the side effects manifested by artemisinin-based medications, including dizziness, nausea, vomiting and loss of appetite.^[12]^

The molecular occurrences upon the activation of artemisinin, generating detrimental reactive oxygen species (ROS), ultimately inducing cell apoptosis.^[13-15]^ Currently, the drug delivery systems are developed to release the drug in response to external ROS stimuli (e.g., H_2_O_2_).^[16-18]^ Although this strategy provides an effective approach for cancer treatment by enhancing permeability and retention (EPR) effect, the clinical translation of these prodrugs has been limited.^[19-21]^ Generally, adverse events associated with toxic side effects and inadequate therapeutic compliance caused by the increased dosing frequency, leading to cancer treatment failure.^[22-24]^ Apparently, inadequate accumulation of drugs within the tumor microenvironment, which consequently undermines the therapeutic efficacy. Additionally, during the circulation process, prodrugs are inevitably taken up by normal tissues, which consequently leads to unfavorable side effects. To our knowledge, no examples of artemisinin-conjugated probes for reducing side effects (AC-Pr-RSE) in vitro/in vivo analyses have been reported so far. Therefore, it is imperative to achieve intratumoral accumulation of prodrugs and enhance the cell killing effect of activated artemisinin.

Artemisinin-derived fluorescence tracers for localization ROS (e.g., H_2_O_2_) imaging, especially in Golgi apparatus are rare. This strategy has the potential to mediate the action of antioxidant defenses and artemisinin cytotoxicity. To address this issue, a H_2_O_2_-triggered theranostic probe (YCF-T) was developed for overcoming the stability dilemma based on AC-Pr-RSE mechanism. We propose that the safe and spacious harbor (nitrogen mustard)-N-methyliminodiacetic acid (MIDA) boronates as a specific H_2_O_2_ recognition group for overcoming the stability in the blood circulation. Moreover, utilized as a targeting moiety, the phenylsulfonamide is employed for targeting the Golgi apparatus in cancer cells, which allows covalent attachment to the theranostic probe. Notably, it is important to note that artemisinin and its derivatives can induce oxidative stress, thereby causing cancer-cell death in the Golgi apparatus. To the best of our knowledge, this is the first example of using naphthalimide fluorophore-mediated Golgi accumulation of H_2_O_2_ based on AC-Pr-RSE mechanism in situ.

**Scheme 1.**
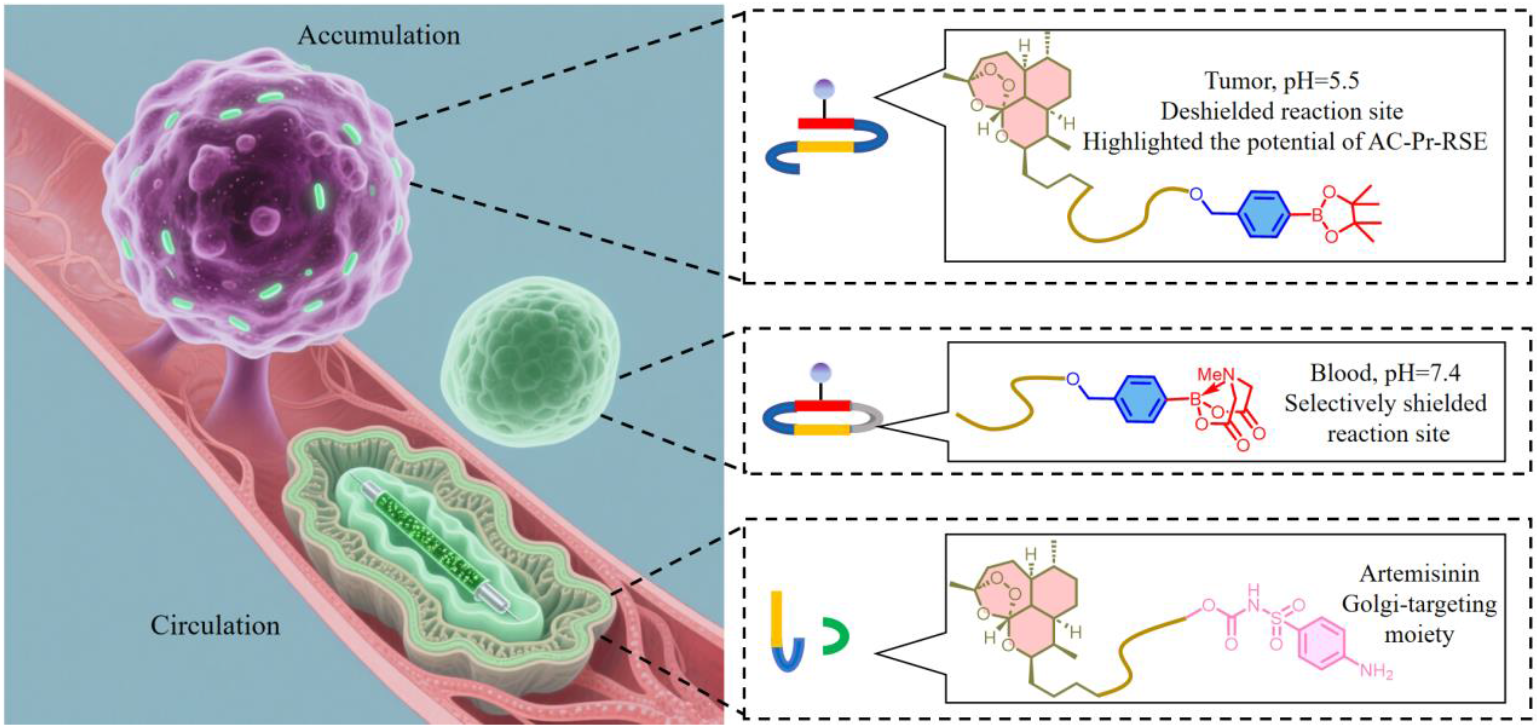
Schematic representation of the reaction of probe with Golgi H_2_O_2_.

## 2. Experimental section

### 2.1 Materials and Instruments

Artemisinin was purchased from Sigma Aldrich. All other chemicals were purchased from major commercial suppliers without further purification unless otherwise noted. The used Silica gel (Silicycle, 300–400 mesh) for column chromatography was prepared from the Alfa Aesar (Tianjin). ^1^H and ^13^C NMR for the target compounds were performed on a Bruker Advance-500 spectrometer, using tetramethylsilane (TMS) as an internal standard. Electrospray ionization mass spectra (ESI-MS) data were measured with Bruker Esquire 3000. Fluorescence imaging experiments were performed on a a F-7000 fluorescence spectrophotometer (Hitachi). Living cell imagings were obtained using an Olympus FV1000-MPE multiphoton laser scanning confocal microscope (Japan). All the optical measurements were performed at room temperature. The synthetic methodology of probe (YCF-T) and analytical data were shown in the Supporting Information.

### 2.2 Cell culture

The cells were cultured in DMEM medium (high glucose) supplemented with 10% fetal bovine serum (FBS, GIBCO) and 1% penicillin-streptomycin (penicillin: 10,000 U·mL^−1^, streptomycin: 10,000 U·mL^−1^) at 37 °C in humidified atmosphere containing 20% O_2_ and 5% CO_2_. Upon reaching an approximate confluence of 80%, cells were passaged using 0.25% trypsin. Fluorescent imaging of probe-incubated cells was performed with a Leica TCS SP5X confocal microscope. The cells were seeded at a density of 1×10^5^ cells per well in a glass-bottom culture dish. Upon a 24-hour incubation to ensure cell adhesion, the culture medium was aspirated, and the cells were rinsed with PBS. Subsequently, the cells were subjected to a 1-hour incubation with probes in DMEM at 37 °C under 5% CO_2_, after which they were washed three times with PBS thoroughly. Confocal fluorescence imaging was conducted on a Leica SP8 confocal microscope (Germany) with 488 nm excitation, and the fluorescence emission windows were set at 520–560 nm.

### 2.3 Model mice

All animal experiments were conducted in accordance with the National Institutes of Health guidelines for experimental animal use (China) and were approved by the Animal Ethics Committee of Guizhou Minzu University. Female BALB/c mice at 8 weeks of age were obtained from Western Biomedical Technology Lab Animal Co., Ltd. (Chongqing, China). To eliminate interference with imaging, the animals were kept in an environment with consistent temperature (22 ± 2 °C) and humidity (50 ± 10%) controlled throughout the imaging process.

## 3. Results and Discussion

### 3.1 Screening and optimization of YCF-T-based sensing method for H_2_O_2_ detection

After syntheses, we first assessed the responsive ability of probe after reaction with H_2_O_2_ by fluorescence spectrum in PBS buffer. As illustrated in Figure 1a, the fluorescence of the solution undergoes hardly any alteration under excitation at 485 nm without H_2_O_2_. Owing to the “Targeted shielding” effect, the fluorescence signal of the probe YCF-T underwent marked silencing. Expectedly, upon adding H_2_O_2_ (130 μM), the fluorescence signal of the probe displayed a significant increasing trend, which suggests that the MIDA) boronates group, as a recognition group, has high reactivity toward H_2_O_2_. Subsequently, it could be observed from Figure 1b that the fluorescence signal of probe YCF-T at around 535 nm was enhanced upon the addition of H_2_O_2_ at various concentrations. This indicates that the uncaging and subsequent release of the fluorophor from YCF-T can be achieved over an extended range of H_2_O_2_ concentrations.

**Figure 1.**
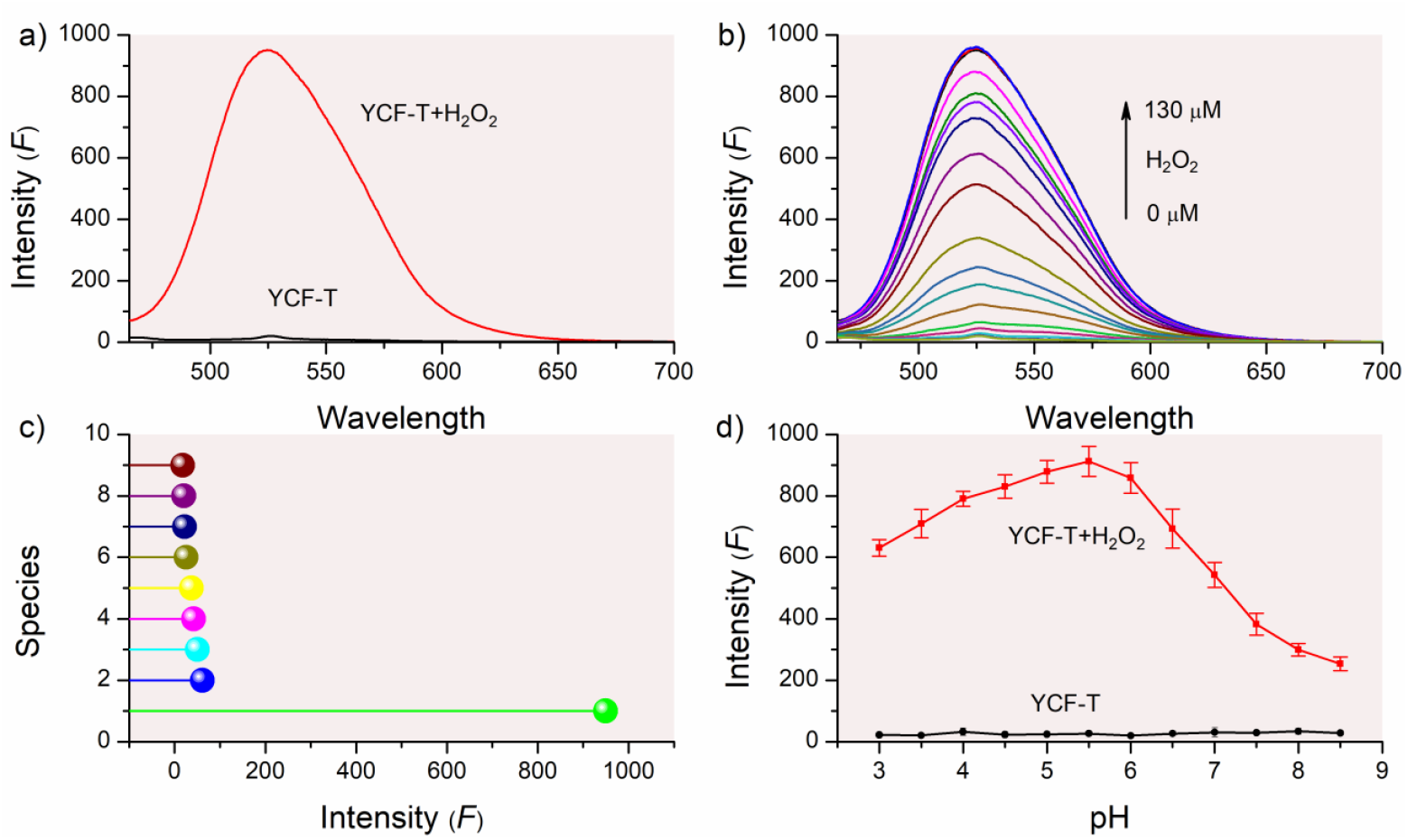
a) Fluorescence emission spectra of YCF-T: Fluorescence emission spectra of YCF-T (1 μM) in phosphate-buffered saline (PBS, pH = 5.5) with or without incubation with H_2_O_2_ (130 μM) at 37 ? for 4 h. Excitation wavelength (λ_em_) = 485 nm. b) Fluorescence emission spectra of YCF-T at different H_2_O_2_ concentrations: Fluorescence emission spectra of YCF-T (1 μM) in the presence of various concentrations of H_2_O_2_, measured in PBS (pH = 5.5). λ_ex_ = 485 nm. c) Fluorescence intensity of YCF-T toward different reactive oxygen species (ROS): Fluorescence intensity of YCF-T (1 μM) at the emission wavelength of 523 nm (λ_em_ = 523 nm) in response to various reactive oxygen species (ROS). d). Effect of pH on the fluorescence spectra of YCF-T: Fluorescence spectra of YCF-T (1 μM) in buffer solution, investigated under varying pH conditions both in the absence and presence of 130 μM H_2_O_2_.

To further investigate the responsive decomposition performance of YCF-T toward H_2_O_2_, various relevant interfering substances were tested and analyzed in buffer solution (20 mM, pH 7.4) under physiological conditions, including hypochlorite (ClO^-^),·OH, and ONOO^−^. As shown in Figure 1c, a striking fluorescence enhancement was observed exclusively with H_2_O_2_, while the fluorescence signal stayed almost steady when other biologically relevant species were added. These experimental results indicate that YCF-T can serve as a specific fluorescent probe for detecting H_2_O_2_, especially when compared to other related analytes under complex physiological and biological conditions. To further evaluate the ROS-induced release behavior, the profiles of YCF-T was carefully monitored at both the acidic tumor microenvironment pH of 5.5 and the physiological pH of 7.4 at 37°C. The pH working range of probe LGM - XL, in both the absence and presence of H_2_O_2_, was investigated by recording fluorescence intensity. Intense fluorescence was observed in acidic media (pH 5.5), indicating that YCF-T could sensitively detect H_2_O_2_ under weakly acidic conditions and holds potential for noninvasive visualization of H_2_O_2_ in vivo. This method takes advantage of the AC-Pr-RSE effect upon the exposure to 130 μM.

Subsequently, we studied the sensitivity of YCF-T towards H_2_O_2_, which involved measuring the limit of detection (LOD) of probe. The plateau of fluorescence intensity was detected after the addition of 9 equivalents of H_2_O_2_ with excitation at 480 nm (Figure 2a). A linear equation F = 12[YCF-T] (μM) - 150, which showed a good linear relationship (R^2^=0.980), was derived as the concentration of YCF-T rose from 0.2 μM to 1.8 μM. The detection limit (LOD) was found to be 1.0 μM based on a signal to noise ratio (S/N= 3), indicating the superior sensitivity of probe towards H_2_O_2_. Additionally, the pseudo-first-order rate constant was found to be 0.115 M^−1^ min^−1^ and the nonlinear curve is shown in Figure S8 (R^2^=0.994). The time-course kinetics of the reaction between YCF-T and H_2_O_2_ was also employed and the results were displayed in Figure 2b. The treatment of YCF-T (1 μM) with 10 equiv of H_2_O_2_ resulted in a remarkable fluorescence enhancement and the balance point after 20 min, indicating artemisinin derivative can induce oxidative stress based on AC-Pr-RSE effect. These results clearly indicated the success of our optimized design for probe in the presence of H_2_O_2_ generated from artemisinin.

**Figure 2.**
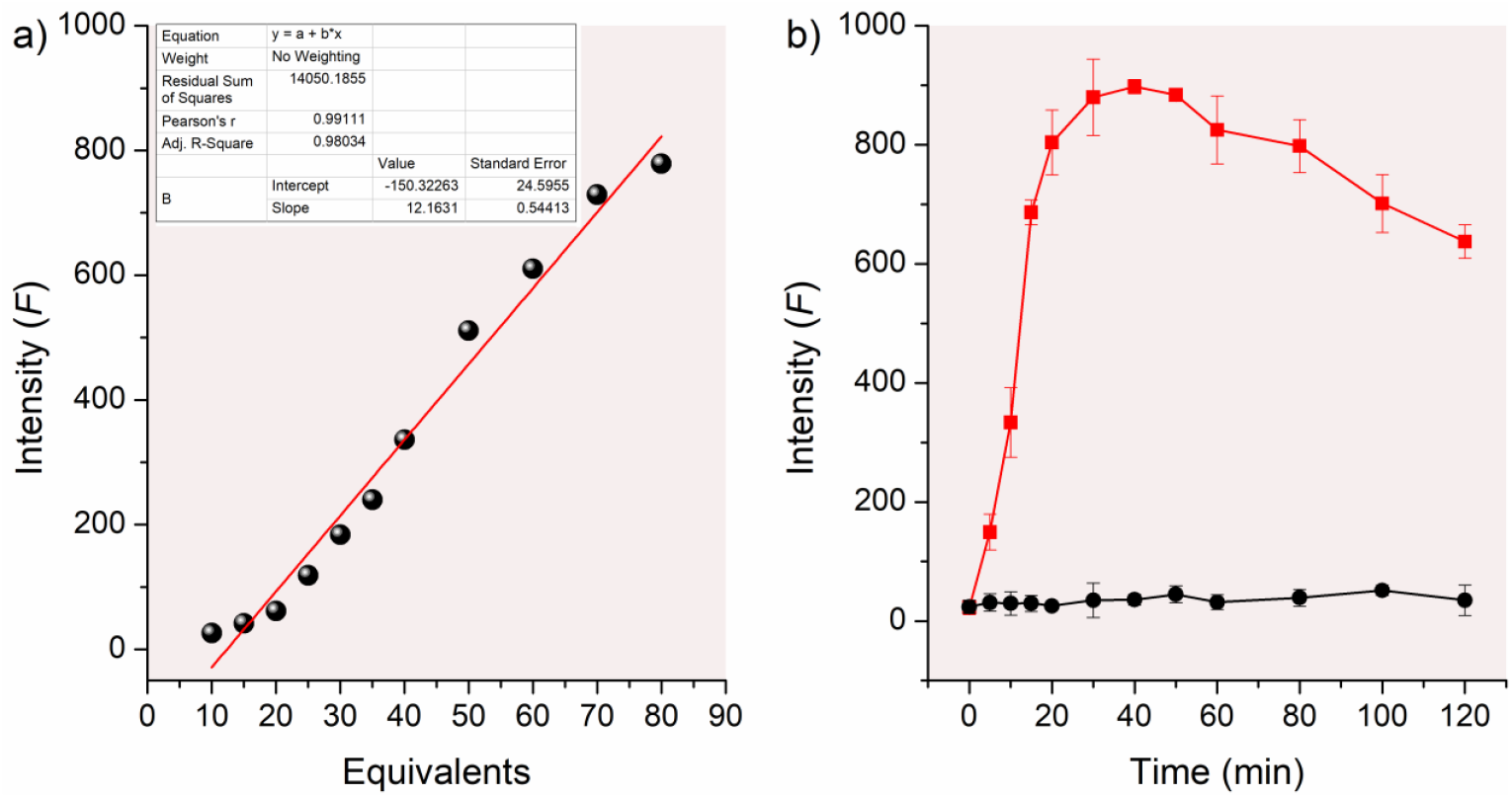
a) Plots of YCF-T fluorescence intensity as a function of H_2_O_2_ concentration: Plots depicting the fluorescence intensity of YCF-T (1 μM) versus the concentration of H_2_O_2_ (tested over a defined concentration range). Excitation wavelength (λ_ex_) = 485 nm; experimental conditions (e.g., buffer type, pH, temperature) were consistent with Figure 1 unless otherwise specified. b) Time-dependent changes in YCF-T fluorescence intensity at pH 5.5: Changes in the fluorescence intensity of YCF-T (1 μM) over different incubation time periods, measured in a buffer system at pH 5.5. Excitation wavelength (λ_ex_) = 485 nm; emission wavelength (λ_em_) was consistent with Figure 1 (e.g., 523 nm) unless otherwise noted.

### 3.2 Sensing mechanism for selective detection of H_2_O_2_ using YCF-T

As indicated by the results compiled in Scheme 2, it is worthwhile to determine whether theranostic probe exhibits similar functionality in live cells. We settled the reaction by mixing and stirring YCF-T with an equivalent amount of H_2_O_2_ for 1 h under the standard condition. In this process, an unstable cyclic intermediate was formed by the spectra of the fluorescence of YCF-T itself and fluorophore, which can make the probe fluoresce at 523 nm. This hydrolysis-triggered substitution-cyclisation-elimination reaction is triggered by the oxidation of the carbon boron bond initiated by the positive charge developed on the nitrogen. Additionally, upon adding H_2_O_2_, a distinct peak at m/z = 623.3 (calcd. 622.33 for [M]^+^) as well as the phenylsulfonamide byproduct ([M + H]^+^, m/z 173.1) were found (Figure S8), which subsequently undergoes rapid intramolecular cyclization, leading to the release of the prodrug and the activation of the fluorescence signal.

**Scheme 2.**
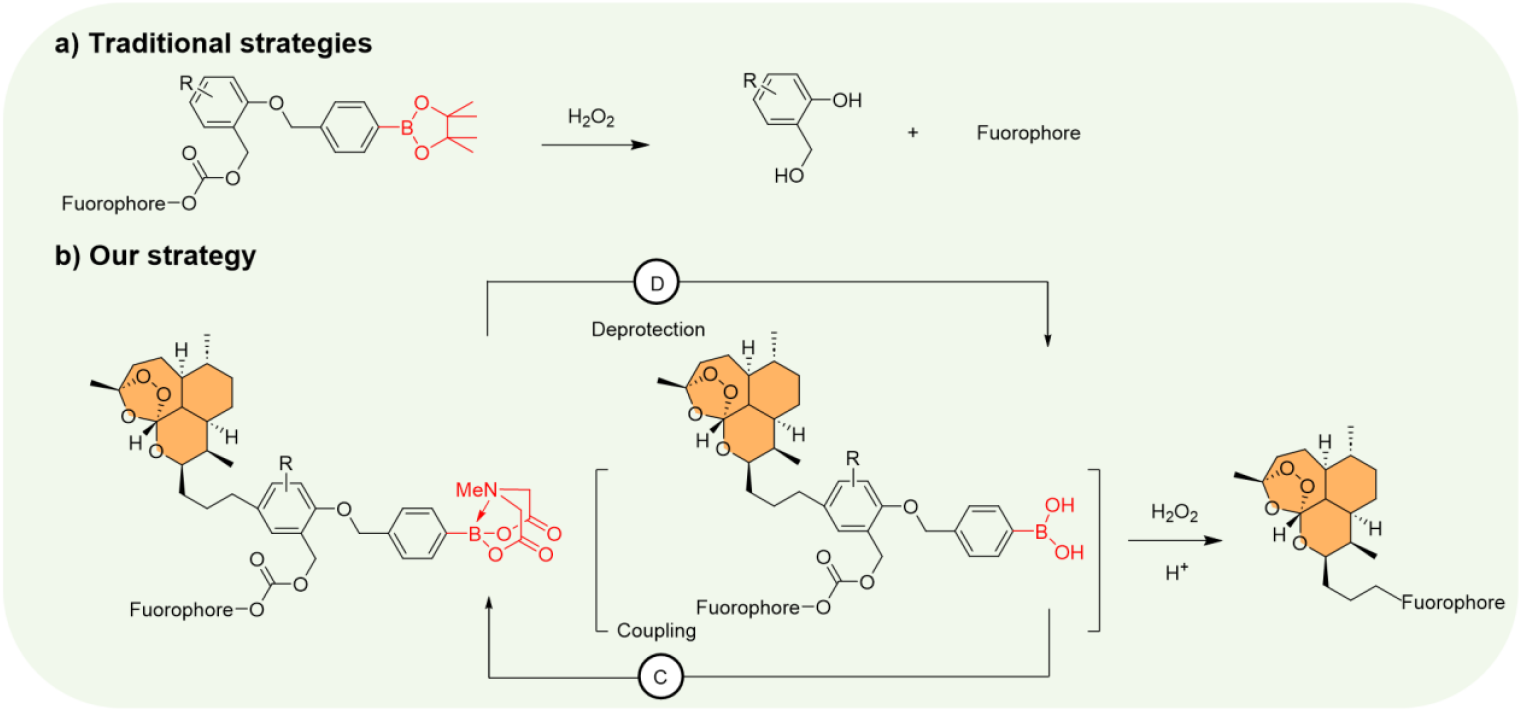
The structure and fluorescence response mechanism of YCF-T.

### 3.3 Cytotoxic Effect of YCF-T

Before applying the probes for cell imaging experiment, we first evaluated the cytotoxicity test toward H_2_O_2_-overexpressing HepG2 cells and normal cells (NIH 3T3 cells). These cells were incubated with various concentrations (0, 0.2, 0.5, 1, 2, 5, 10, 20 and 30 µM) of YCF-T for 48 h. As shown in Figure S9, the cancer cell survival rates remained more than 78% even the concentration of the probe was up to 30 µM after incubation for 24 h according to normal cells, suggesting that YCF-T possesses low cytotoxicity and and can serve as a desirable probe for constructing controlled drug release systems.

### 3.4 Golgi-targeting capability of YCF-T

Subsequently, taking into account the exceptional detection performance of YCF-T, the link between cellular inflammation and H_2_O_2_ expression was further investigated in various types of living tumor cells (HepG2 and RAW264.7 cells). As depicted in Figure 3, we employed a commercially available Golgi probe Golgi-Tracker Red to co-stain with YCF-T (λ_ex_= 485 nm, λ_em_ = 523 nm) before imaging under confocal laser scanning microscopy (CLSM). The merged images indicated that the green fluorescence of the probe overlapped well with the fluorescence of Golgi-Tracker Red in the Golgi apparatus of both RAW264.7 cells and HepG2 cells, with Pearson’s colocalization coefficients of 0.93 and 0.95, respectively. The results indicated the strong affinity of YCF-T to the Golgi apparatus through the synergistic targeting effects of the phenylsulfonamide during various pathological processes.

**Figure 3.**
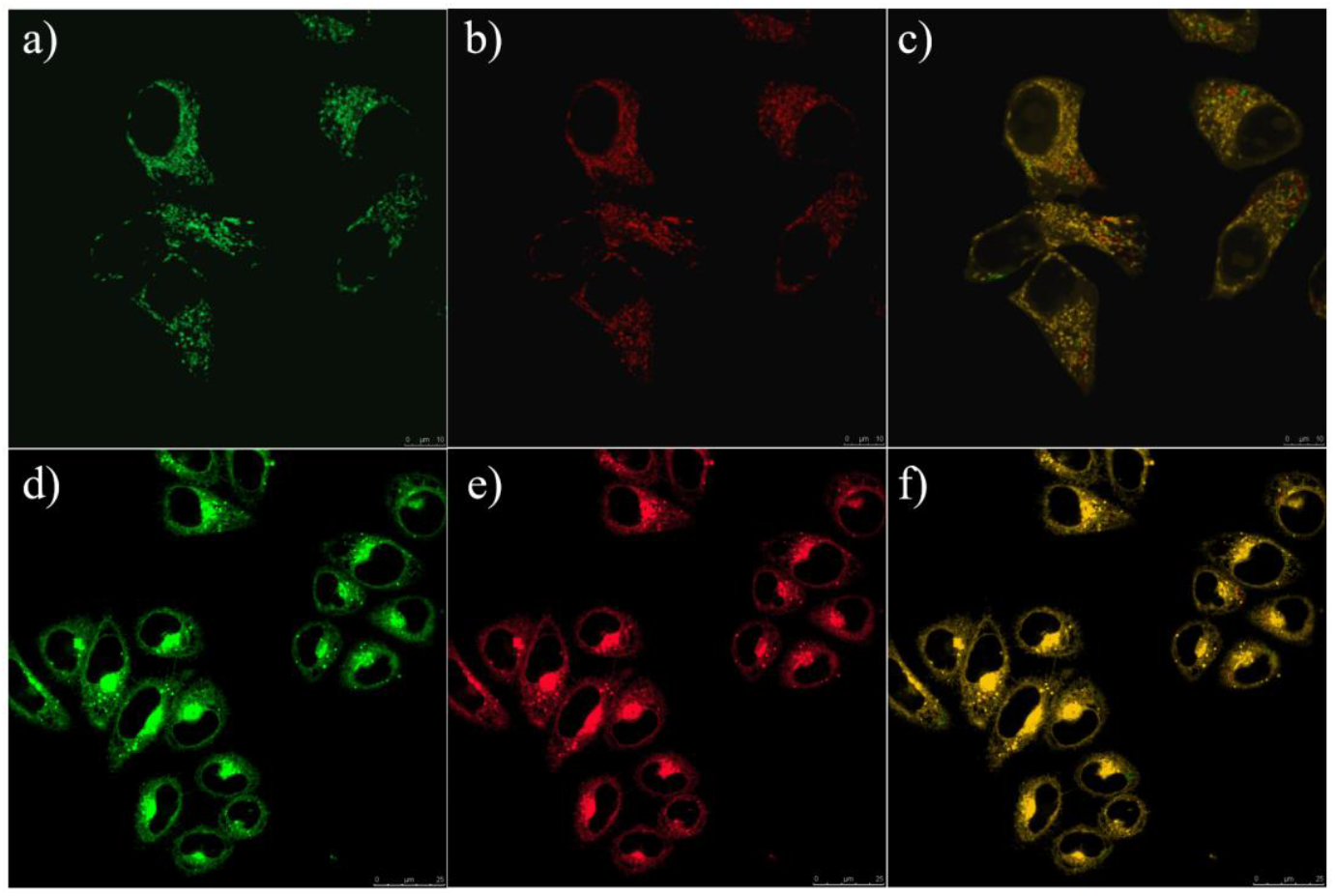
Co-localization cell imaging of YCF-T (1 μM) and commercial dyes in RAW264.7 cells (a, b, c) and HepG2 cells (d, e, f).

To demonstrate its capability for cellular H_2_O_2_ fluorescence imaging, cells were incubated with YCF-T and a reference molecule lacking artemisinin unit (YCF) at 37 ° for 30 min (Figure 4). Encouragingly, green fluorescence emission was observed inside the cells by using a confocal fluorescence microscope (Figure 4a), demonstrating that artemisinin dependent endoperoxide bioactivation take place in Golgi apparatus. A markedly lower fluorescence intensity increase takes place when the cells are pretreated with YCF for 30 min, as is shown in Figure 4b. These findings demonstrate that artemisinin triggered H_2_O_2_ synthesis and oxidative stress.

**Figure 4.**
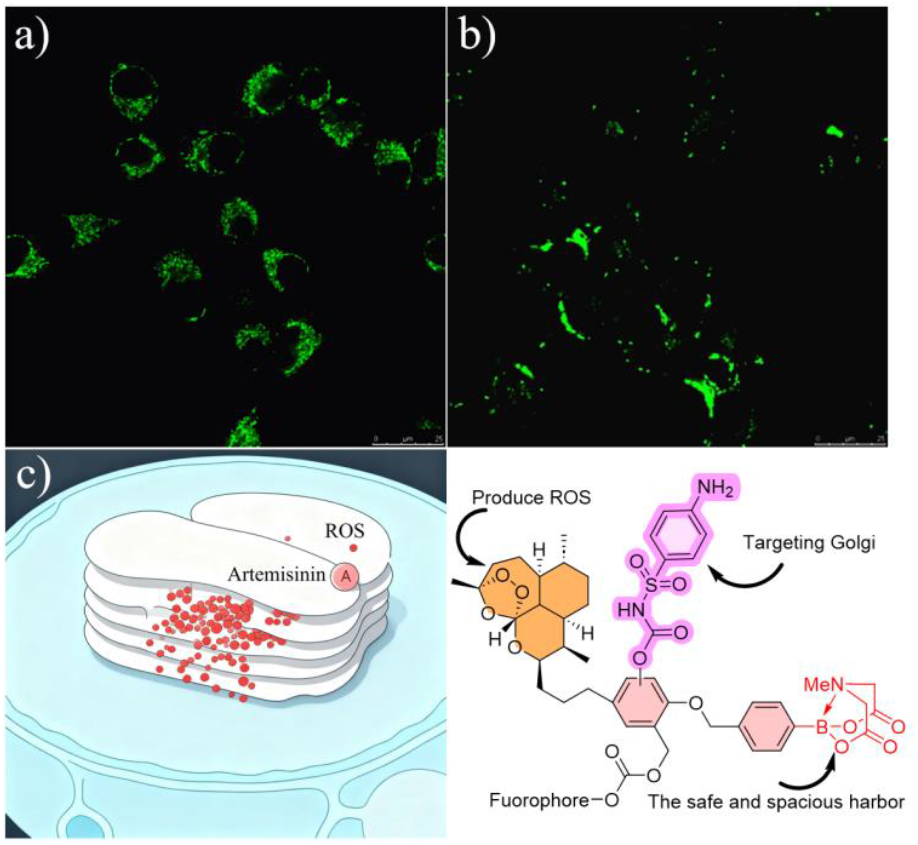
Fluorescence of cells treated with YCF-T (a) and YCF (b, without artemisinin) for 24 h.

### 3.5 The recognition of probe YCF-T to H_2_O_2_ in cellula

After exploring YCF-T function, we evaluated the physiological activity of artemisinin on the change of H_2_O_2_ concentration in the Golgi apparatus. The effects of AC-Pr-RSE on the staining of YCF-T in the Golgi of HepG2 cells were investigated, and the results were shown in Figure 5a. Next, the brighter fluorescence of YCF-T in the green channel (530–580 nm) increased significantly upon the addition of Monensin (Mone, 2 mM), which was reported as the specific expression vector for endogenous H_2_O_2_ within the Golgi apparatus (Figure 5b). However, when cells were exposed to the ROS-scavenger N-acetyl cysteine (NAC, 2 mM), the H_2_O_2_ levels in the NAC-stimulated cells decreased by approximately 2.5-fold (Figure 5c). These data displayed that the level of Golgi H_2_O_2_ is obviously increased in the Golgi oxidative stress process based on AC-Pr-RSE mechanism.

**Figure 5.**
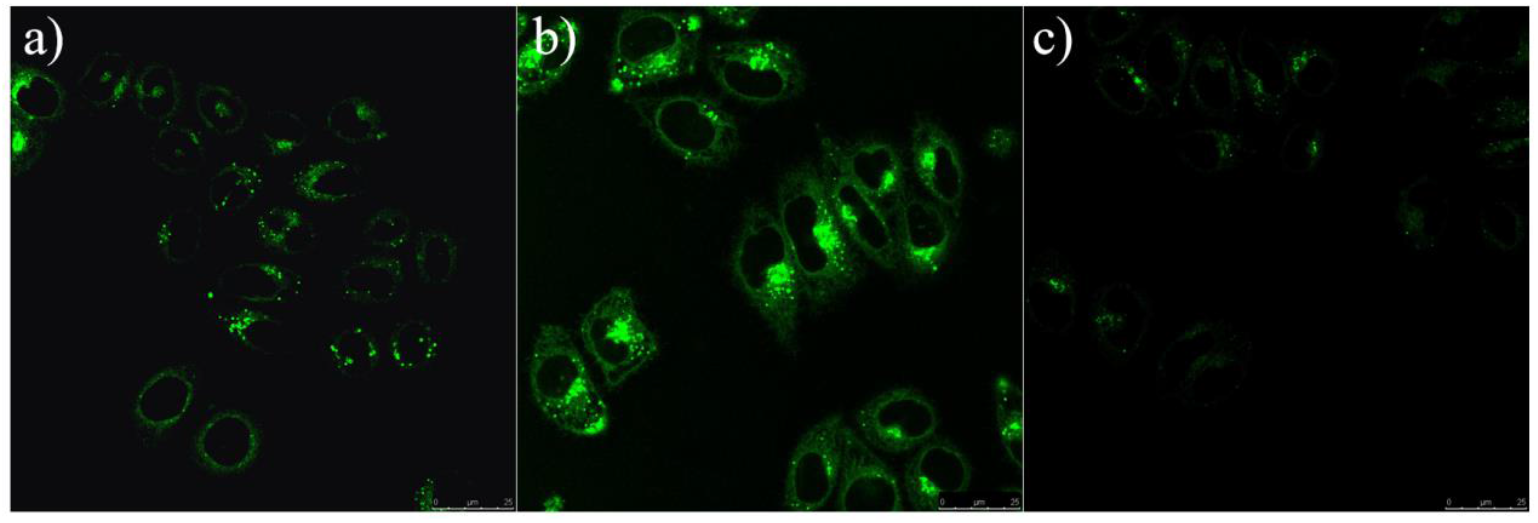
The cell imaging of YCF-T (1 μM) in HepG2 cells (a). Fluorescence imaging of YCF-T after cells were treated with different stimulants. (b) NAC cells: The cells were incubated with NAC (20 mM). (c) Monensin cells: The cells were incubated with NAC and then monensin (10 mM) was added.

Encouraged by the findings above, the time-dependent imaging assay enabled continuous monitoring of H_2_O_2_ activity in live cells, and this temporal investigation was extended to HepG2 cell. As shown in Figure 6, incubation of cells with YCF-T led to a progressive enhancement of fluorescence over time, which plateaued at 20 min and maintained stability for the following 40 min, but a substantial reduction in fluorescence signal was observed after 4 h, suggesting the gradual translocation of the precipitated fluorochrome YCF-T out of live cells. These data clearly demonstrate that the outstanding signal-stability of YCF-T with maximizing contrast and high sensitivity.

**Figure 6.**
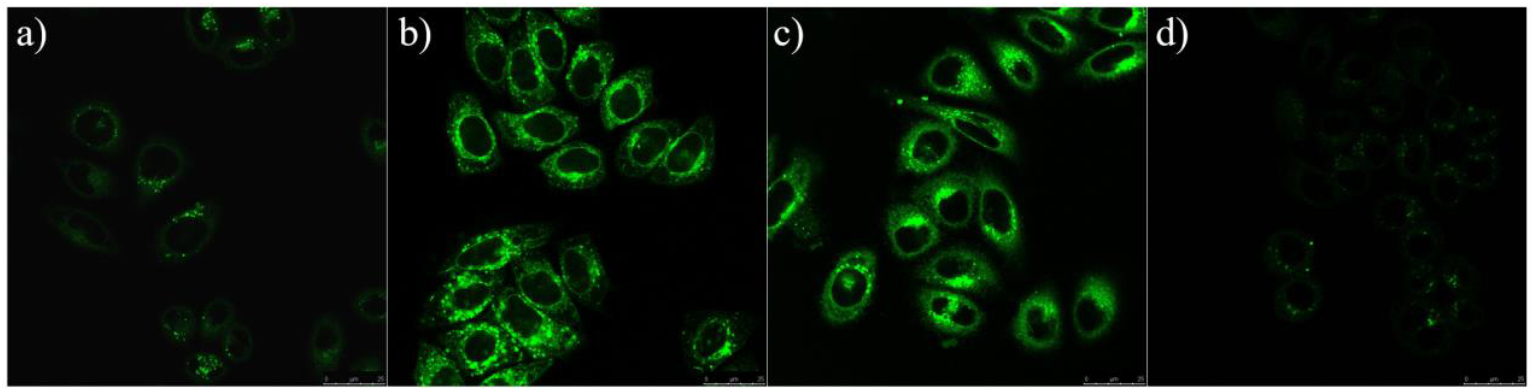
Confocal images of HepG2 cells. Cells were treated with 1 μ? YCF-T for (a) 10 min, (b) 20 min, (c) 40 min and (d) 4 h.

### 3.6 Tissue imaging of H_2_O_2_ in inflamed mouse model

To assess whether YCF-T is detectable in vivo within the Golgi of macrophages in a mouse model of inflammation, 100 μL of LPS (1 mg/mL) was delivered subcutaneously into the left hind paws of the experimental mice to induce inflammation (Figure 7). One day after the previous LPS administration, YCF-T was delivered via intravenous injection to the experimental mice. Subsequently, the left hind paw skin was excised and sectioned one hour post-injection. As shown in Figure 7b, strong fluorescence signal from YCF-T was observed in LPS-stimulated macrophages located in the inflammatory tissues, and these signals were visualized in green channel. However, the signal of the probe and the macrophage marker were largely absent in the normal tissues. (Figure 7c and S10). Taken together, YCF-T successfully stained inflamed mouse macrophages via intravenous injection, indicating higher H_2_O_2_ in their Golgi during inflammation.

**Figure 7.**
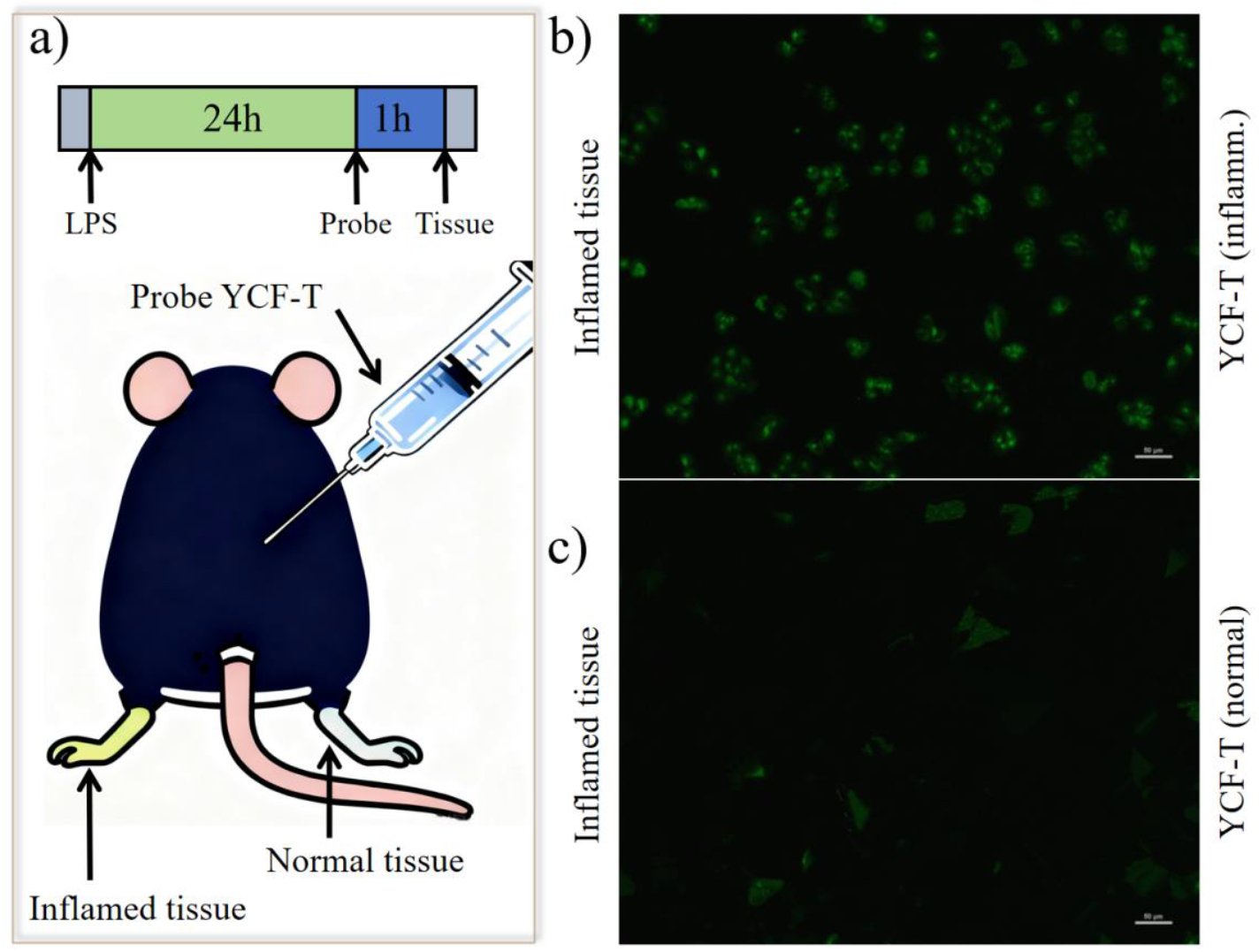
Measurement of LPS-Mediated H_2_O_2_ generation in inflammatory tissues via YCF-T. (a) Mice received a subcutaneous (s.c.) administration of 100 μL LPS (1 mg/mL) into the left hind paw to induce inflammation. One day after this LPS administration, 100 μL of 1 mM YCF-T was administered intravenously (i.v.), and the paw skin was subsequently excised and sectioned 1 hour later. (b) (c) Fluorescence images of probe in the inflamed tissue. Probe fluorescence, green.

## 4. Conclusions

In summary, this work presents the first report of a theranostic platform constructed by integrating a cytotoxic artemisinin derivative, N-methyliminodiacetic acid (MIDA) boronates, and a naphthalimide fluorophore, which is capable of detecting H_2_O_2_ in the Golgi apparatus. Through comparative analysis of fluorescence localization signals with organelle-specific dyes, we confirmed that the Golgi apparatus acts as the primary accumulation site for this artemisinin-based probe. Functionally, such targeted accumulation was demonstrated to efficiently upregulate intracellular reactive oxygen species (ROS) levels and induce cellular apoptosis. These findings not only advance our mechanistic understanding of artemisinin-conjugated phototheranostic probes for side effect reduction (AC-Pr-RSE) but also, more importantly, provide critical insights for the rational optimization of theranostic probe design. This, in turn, is expected to enhance their biological activity under oxidative stress conditions, thereby contributing to the improvement of therapeutic efficacy in relevant applications.

## Acknowledgment

We would like to acknowledge the financial support of the National Natural Science Foundation of China (NSFC) projects (22267007), the Guizhou Provincial Basic Research Program (Natural Science) (No. Qian Ke He Basic - [2025] Youth 312), the Longyuan Youth Innovation and Entrepreneurship Talent Project of Gansu Province (2023LQGR15).

## Notes

### Competing Interest Statement

The authors have declared no competing interest.

